# Interocular suppression weakens binocular disparity integration in the dorsal visual stream

**DOI:** 10.64898/2026.09.11.751085

**Authors:** Rong Jiang, Ming Meng

**Affiliations:** State Key Laboratory of Eye Health, Eye Hospital, Wenzhou Medical University, Wenzhou, 325027, China; Department of Neurobiology, The University of Alabama at Birmingham, Birmingham, USA; School of Psychology, South China Normal University, Guangzhou, China

**Keywords:** stereopsis, binocular rivalry, interocular suppression, binocular disparity, fMRI, multivariate pattern analysis

## Abstract

A unified percept depends on the coordinated interplay of sensory integration and suppression, yet how this interaction is implemented across cortical representations remains unclear. Using binocular vision as a tractable model system, we combined fMRI with multivariate analyses to test how interocular suppression reshapes binocular disparity integration across human visual pathways. Local binocular rivalry weakened near–far disparity decoding, with disruption increasing along the dorsal pathway from V1 to V3A and intraparietal sulcus (IPS), but no reliable loss in ventral regions. In V3A, anticorrelated random-dot stereograms showed an inverted representational relationship relative to correlated stimuli, and this structure was abolished under suppression, indicating disrupted non-canonical disparity integration. Informational connectivity further revealed that suppression selectively weakened V3A-centered coordination while largely sparing coupling among early visual areas. Together, these findings identify higher-level dorsal regions as a key substrate for coordinating integrative and suppressive computations during unified percept formation.

## Introduction

How the brain constructs a coherent percept from multiple sensory inputs is a central question in neuroscience (Blake, 2022; Blake & Wilson, 2011; Stein & Stanford, 2008). Achieving this depends on the interplay of two complementary computations: the integration of compatible signals and the suppression of conflicting ones (Carandini & Heeger, 2012; Desimone & Duncan, 1995; Friston & Kiebel, 2009). Binocular vision offers a particularly tractable framework for investigating this interplay, as compatible signals from the two eyes can be integrated to support stereoscopic depth perception (Henriksen et al., 2016; Nityananda & Read, 2017), whereas conflicting inputs can trigger interocular suppression, resulting in binocular rivalry (Blake, 2022; Blake & Wilson, 2011; Tong et al., 2006). Recent psychophysical findings suggest that binocular integration and suppression are intrinsically linked (Blake, 2022; Blake & Wilson, 2011; Jiang et al., 2025; Jiang & Meng, 2023, 2025). Yet how their interaction is coordinated across different levels of cortical processing remains unclear.

Extensive prior work has identified the dorsal visual stream as a key pathway for both binocular integration and interocular suppression. Inputs from the two eyes first converge in V1, providing a common cortical entry point for both processes. In binocular integration, V1 neurons begin to encode disparity information (Henriksen et al., 2016; Ohzawa et al., 1990), and disparity representations are further elaborated across visual cortex, with dorsal regions such as V3A and intraparietal sulcus (IPS) playing especially important roles in behaviorally relevant depth processing (Georgieva et al., 2009; Kennedy et al., 2023; Rosenberg et al., 2023; Tsao et al., 2003). In interocular suppression, neural signatures likewise extend from early visual cortex to parietal regions (Haynes et al., 2005; Lunghi et al., 2015; Polonsky et al., 2000; Tong & Engel, 2001; Wunderlich et al., 2005; Zaretskaya et al., 2010), with recent high-resolution 7T fMRI evidence suggesting that V1 contributes to resolving local interocular conflict, whereas IPS supports the broader spatial coordination of perceptual dominance (Qian et al., 2026). The established importance of the dorsal visual stream for both binocular integration and interocular suppression therefore suggests that dorsal-stream regions may provide a key cortical substrate for the interaction between these two processes.

To test this hypothesis, we used functional magnetic resonance imaging (fMRI) to examine how interocular suppression reshapes disparity processing across dorsal visual areas V1, V2, V3, V3A, and IPS, with hV4 and VO included as ventral comparison regions to assess pathway selectivity (Fig. 1B). Random-dot stereograms (RDS) carrying near–far disparity were presented in a peripheral annulus, while the binocular state of the central field was independently manipulated with binocularly matched or mismatched inputs to induce fusion or rivalry (Fig. 1A). Multivariate decoding was then used to quantify disparity information directly from neural activity patterns (Ng et al., 2021; Preston et al., 2008), enabling us to test how interocular suppression reshapes neural representations of binocular disparity without relying on explicit reports of stereoscopic percepts. This decoding-based, non-report approach also enabled the inclusion of anticorrelated RDS, which typically fail to produce stable stereoscopic percepts despite engaging widespread disparity-processing mechanisms (Aoki et al., 2017; Zhaoping & Ackermann, 2018), thereby extending our investigation to non-canonical disparity processing.

**Figure 1.**
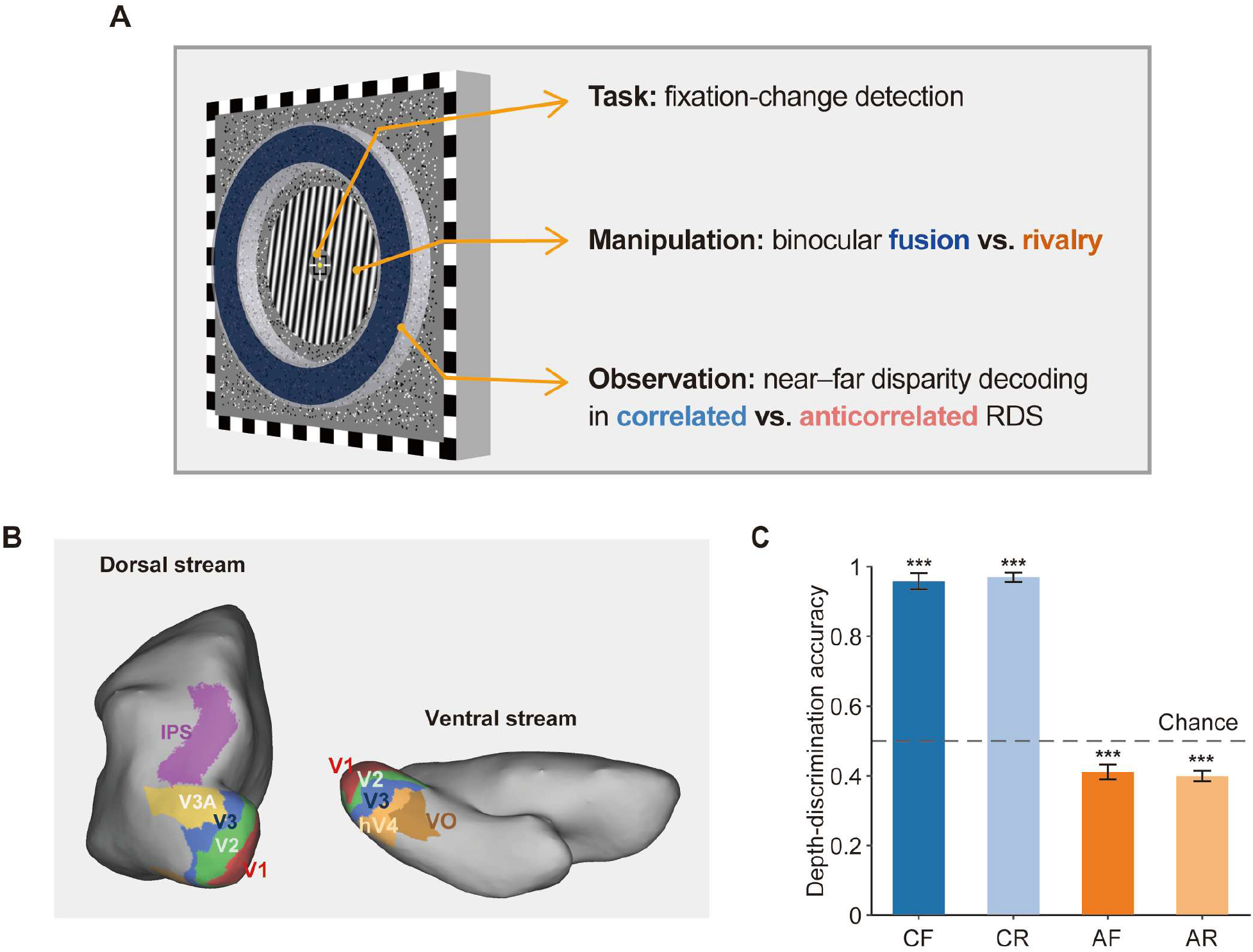
Experimental design and regions of interest. **(A)** During fMRI, participants maintained central fixation and performed a fixation-shape change detection task. The central field (eccentricity 0.4°–2°) contained binocularly matched images (fusion) or mismatched images (rivalry/suppression), while near–far disparity was carried by correlated or anticorrelated random-dot stereograms (RDS) in a peripheral annulus (eccentricity 2.5°–3.5°). **(B)** Regions of interest (ROIs) for decoding analyses. **(C)** Stereoscopic discrimination in a separate psychophysical test. *** denotes *p* < 0.001. CF, correlated fusion; CR, correlated rivalry; AF, anticorrelated fusion; AR, anticorrelated rivalry. Error bars indicate ±SE across participants. *Abbreviations:* IPS, intraparietal sulcus; RDS, random-dot stereograms; ROI, region of interest.

## Results

### Psychophysical validation and matched task engagement

We first used a separate psychophysical test to assess stereoscopic discrimination under the same stimulus parameters as those used in the scanner (±3, ±9, and ±15 arcmin; Fig. 1C). For correlated random-dot stereograms, near–far discrimination was near ceiling under both fusion and rivalry (CF: 0.96 ± 0.07; CR: 0.97 ± 0.04; mean ± SD) and was reliably above chance in both conditions (both *ps* < 0.001). In contrast, anticorrelated stereograms elicited a weak reversal bias, yielding performance reliably below chance (AF: 0.41 ± 0.06, *t*(7) = −4.17, *p* = 0.004, Cohen’s *d* = −1.48; AR: 0.40 ± 0.04, *t*(7) = −6.82, *p* < 0.001, *d* = −2.41). Rivalry did not significantly alter discrimination performance for either correlated or anticorrelated stimuli (both *ps* = 0.838), possibly reflecting reduced behavioral sensitivity to additional rivalry-related effects under ceiling-level correlated performance and weak anticorrelated reversal.

During fMRI scanning, participants performed a central fixation-change detection task. Detection accuracy did not differ across conditions (CF: 0.65 ± 0.19; CR: 0.63 ± 0.21; AF: 0.65 ± 0.21; AR: 0.64 ± 0.20; *F*(3,21) = 1.537, *p* = 0.234, *η*p^2^ = 0.180), indicating comparable task engagement across conditions.

### Suppressive context weakens disparity representations in the dorsal visual stream

We then asked how task-irrelevant interocular suppression affects disparity representations across the cortical hierarchy without relying on explicit perceptual reports (Fig. 2A). Under correlated fusion (CF), near–far disparity information was decodable above chance in all early visual ROIs (V1–V3), both dorsal ROIs (V3A and IPS), and both ventral ROIs (hV4 and VO) (all *ps* ≤ 0.041), with the highest decoding accuracy in V3A (0.67 ± 0.07, *t*(7) = 6.86, *p* < 0.001, *d* = 2.42). Under correlated rivalry (CR), decoding accuracy remained significantly above chance only in V3 (*p* = 0.043) and V3A (*p* = 0.002). For anticorrelated conditions (AF and AR), decoding accuracy did not significantly differ from chance in any ROI (all *ps* ≥ 0.100).

**Figure 2.**
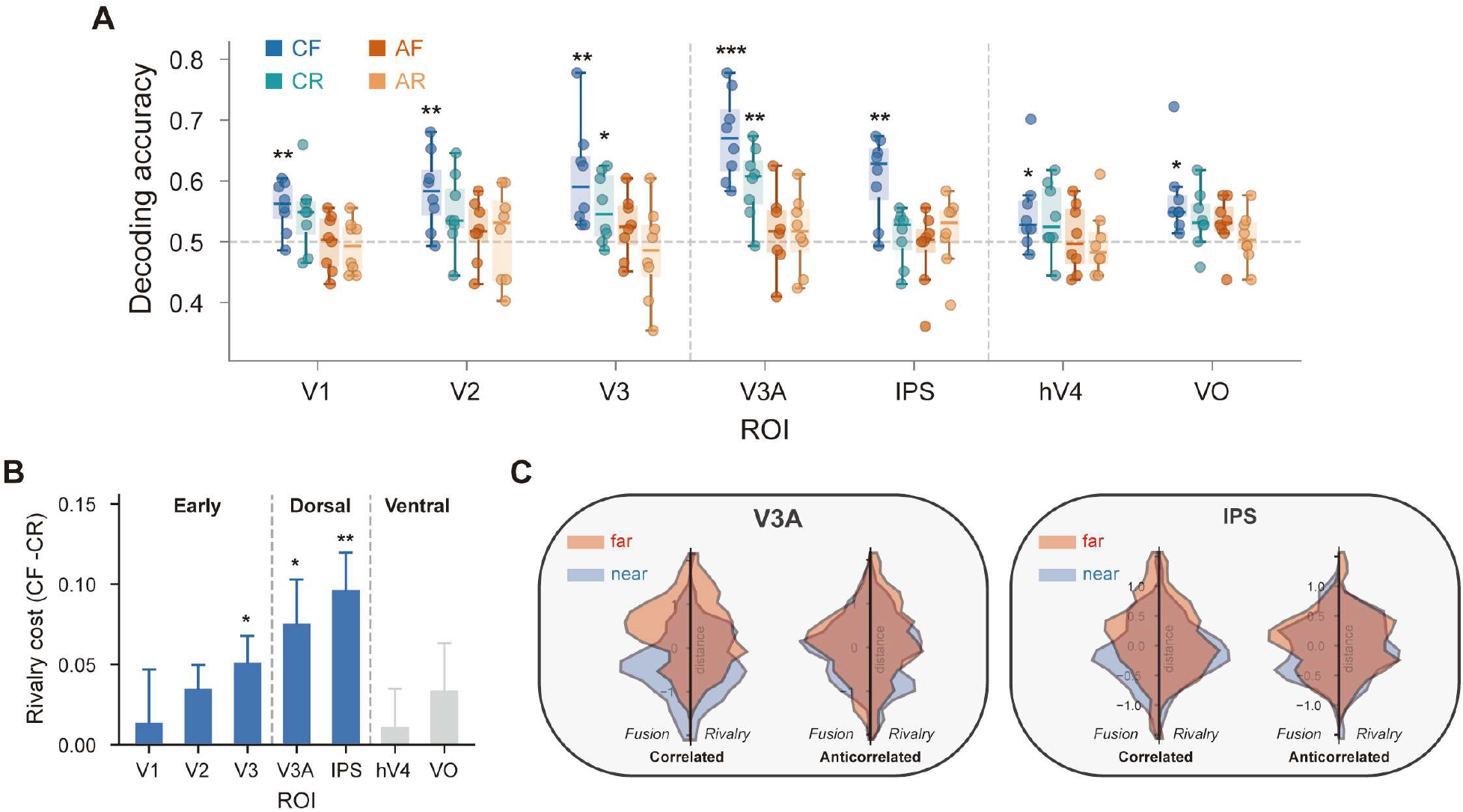
Suppressive context weakens disparity representations across visual cortex. **(A)** Within-condition near–far decoding accuracy across ROIs for CF, CR, AF, and AR. **(B)** Rivalry cost for correlated stimuli (CF − CR). A linear mixed-effects model showed that rivalry cost increased from V1 to IPS along the dorsal visual stream (slope = 0.020, SE = 0.007, *p* = 0.003). **(C)** Block-wise decision values from a representative participant in V3A and IPS. Positive values indicate “far” classifier decisions and negative values indicate “near” decisions. Error bars indicate ±SE across participants. *, **, *** denote *p* < 0.05, 0.01, and 0.001. Abbreviations: CF, correlated fusion; CR, correlated rivalry; AF, anticorrelated fusion; AR, anticorrelated rivalry; ROI, region of interest; SE, standard error.

Direct CF–CR comparisons showed that rivalry weakened correlated disparity representations (Fig. 2B). This rivalry cost was significant in V3 (*p* = 0.027, *d* = 1.09), V3A (*p* = 0.014, *d* = 0.98), and IPS (*p* = 0.004, *d* = 1.47), with a trend in V2 (*p* = 0.053, *d* = 0.82), but was not significant in V1, hV4, or VO (all *ps* ≥ 0.286). Consistent with a hierarchical gradient, a linear mixed-effects model (random intercepts for subjects) showed that the CF–CR rivalry cost increased along the dorsal visual stream from V1 to IPS (slope = 0.020, SE = 0.007, *p* = 0.003).

Fig. 2C illustrates this reduced representational separability in a representative participant, showing the distributions of block-wise decision values for near and far blocks in V3A and IPS across the four conditions. In V3A, decision values for near and far blocks were clearly separated in CF, showed reduced separation in CR, and largely overlapped in AF and AR. In IPS, near–far separation was evident only in CF, with substantial overlap in CR, AF, and AR.

### Cross-condition generalization reveals preserved correlated codes and abolished anticorrelated inverted representations

The reduction in decoding accuracy under suppressive context could reflect weaker disparity signals, a change in representational format, or both. To further probe this issue, we assessed cross-condition generalization across dorsal-stream ROIs using correlated fusion (CF)—the most canonical and interpretable condition—as the anchor for comparison (Fig. 3A). Split-half decoding within CF was robustly above chance in all examined ROIs (all *ps* ≤ 0.003), establishing that the correlated near–far code under fusion was reliable within participants.

**Figure 3.**
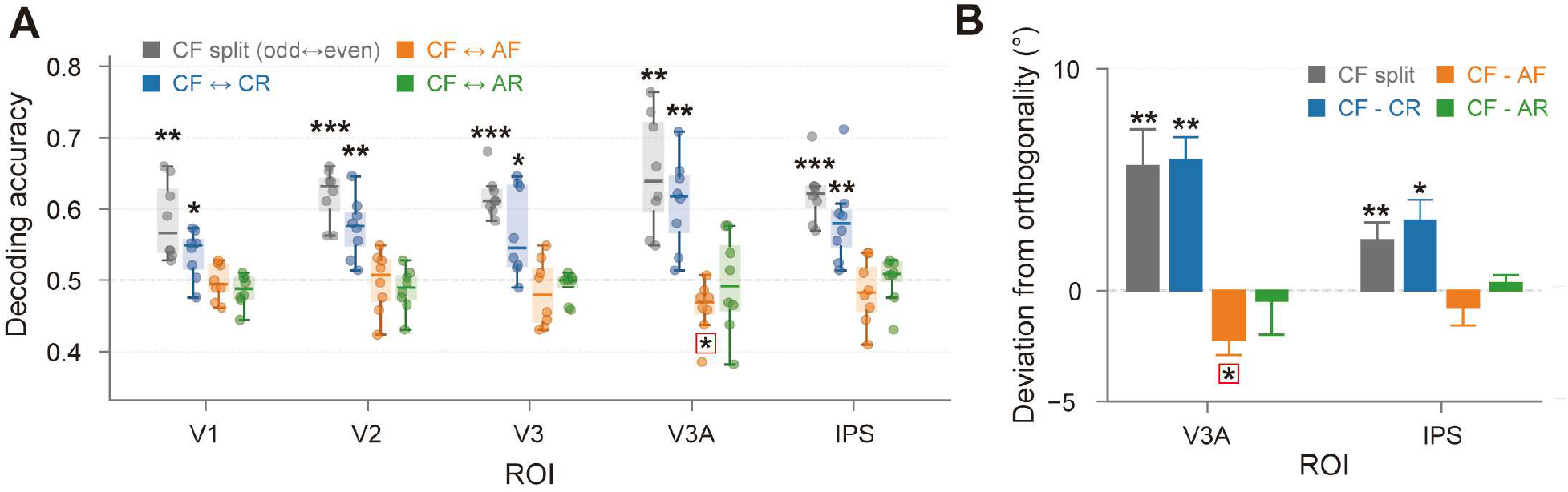
Cross-condition generalization reveals preserved correlated codes and a disrupted anticorrelated disparity mapping in V3A. **(A)** Bidirectional cross-decoding using correlated fusion (CF) as a reference. **(B)** Decision-axis geometry (90° − θ): positive values indicate aligned coding; negative values indicate opponent mapping. Error bars indicate ±SE across participants. *, **, *** denote *p* < 0.05, 0.01, and 0.001. *Abbreviations:* CF, correlated fusion; CR, correlated rivalry; AF, anticorrelated fusion; AR, anticorrelated rivalry; ROI, region of interest; SE, standard error.

Critically, bidirectional cross-decoding between CF and CR (CF↔CR) remained significantly above chance in all examined ROIs (all *ps* ≤ 0.037). Thus, despite the reduced decoding magnitude under rivalry, suppressive context did not abolish the underlying representational format of correlated disparity information.

For anticorrelated conditions, bidirectional cross-decoding with CF was generally at chance in V1, V2, V3, and IPS (all *ps* ≥ 0.324). Notably, in V3A, CF↔AF cross-decoding was significantly below chance (*p* = 0.039, *d* = −1.06), indicating an opponent relationship in which near–far labels tended to reverse between correlated and anticorrelated RDS under fusion. This inverted code disappeared when rivalry was added: CF↔AR in V3A did not significantly differ from chance (*p* = 1.000, *d* = −0.08).

Decision-axis angle analyses converged with the cross-decoding results (Fig. 3B). We quantified axis similarity as the deviation from orthogonality (90° − θ), where positive values indicate aligned coding directions and negative values indicate an opponent mapping. In both V3A and IPS, the angle deviations for the CF split-half and CF vs. CR comparisons were significantly greater than zero (all *ps* ≤ 0.013). By contrast, only in V3A was the angle deviation for CF vs. AF significantly below zero (*p* = 0.031, *d* = −1.13), whereas the deviation for CF vs. AR did not differ from zero (*p* = 0.775, *d* = −0.11). In IPS, the angle deviations for both CF vs. AF and CF vs. AR did not differ significantly from zero (both *ps* ≥ 0.405).

These results suggest that suppressive context largely preserves the representational format of correlated disparity information, but disrupts the non-canonical opponent mapping observed for anticorrelated disparity in V3A.

### Interocular suppression selectively weakens V3A-centered coordination of disparity information

We next asked whether suppressive context alters not only local disparity representations, but also the coordination of disparity information across regions. Within each condition, we performed near–far decoding separately for each region and extracted, for each block, the distance of the activation pattern from the classifier decision boundary. Informational connectivity was then quantified as the Fisher *z*-transformed correlation between these block-wise distance series for selected ROI pairs (Fig. 4).

**Figure 4.**
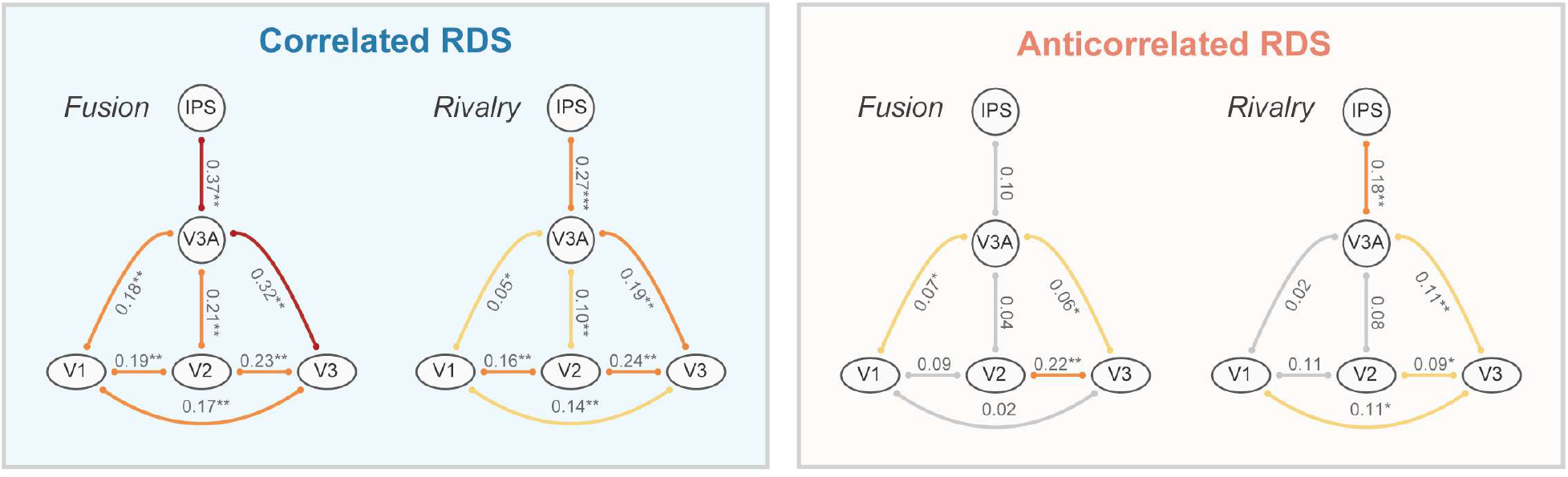
Interocular suppression selectively weakens V3A-centered informational connectivity. Values denote Fisher *z*-transformed correlations between cross-validated decision-value series for selected ROI pairs (within early visual cortex and V3A-centered links). Non-significant connectivity is shown in gray. Significant connectivity is color-coded by effect magnitude: *z* < 0.15, light orange; 0.15 ≤ *z* < 0.3, strong orange; *z* ≥ 0.3, deep red. For correlated stimuli, mean connectivity in the V3A-centered network was lower in rivalry than in fusion (CF > CR), whereas mean connectivity within early visual cortex did not differ between conditions. Anticorrelated conditions showed weaker and more heterogeneous coordination across ROI pairs. *, **, *** denote *p* < 0.05, 0.01, and 0.001. *Abbreviations:* CF, correlated fusion; CR, correlated rivalry; AF, anticorrelated fusion; AR, anticorrelated rivalry; ROI, region of interest.

Under correlated fusion (CF), informational connectivity was significantly greater than zero for all tested ROI pairs, including links within early visual cortex (V1–V2, V2–V3, and V1–V3) and all V3A-centered links (V3A–V1, V3A–V2, V3A–V3, and V3A–IPS; all *ps* ≤ 0.006). Under correlated rivalry (CR), informational connectivity values were numerically reduced for V3A-centered links, but all tested ROI pairs still remained significantly above zero (all *ps* ≤ 0.040). These results indicate that suppressive context did not abolish coordinated disparity-related fluctuations across regions.

Direct CF–CR contrasts, however, revealed a selective reduction in large-scale coordination centered on V3A. At the network level, mean informational connectivity was significantly lower in CR than CF for the V3A-centered network (*p* = 0.024, *d* = 1.24), whereas mean connectivity within early visual cortex did not differ between conditions (*p* = 1.000, *d* = 0.10). Thus, interocular suppression selectively weakened disparity-information coordination in the V3A-centered network while sparing coordination within early visual cortex.

In anticorrelated conditions, informational connectivity was generally weaker and less consistent than in correlated conditions, with reliable positive coupling observed only in a subset of ROI pairs (see Table S6 for details). Moreover, direct AF–AR contrasts revealed no reliable differences at either the ROI-pair level or the network level (all *ps* ≥ 0.148), suggesting that rivalry did not systematically modulate the already weak and poorly organized coordination of disparity information under anticorrelation.

## Discussion

### IPS and the cross-space coordination of binocular integration and interocular suppression

Binocular vision provides a useful model for understanding how integrative and suppressive computations interact: signals from the two eyes can be integrated to support stereopsis or suppressed to give rise to rivalry. Increasing evidence suggests that these processes are closely linked components of a common binocular system (Jiang & Meng, 2023, 2025; Riesen et al., 2019; Wilson, 2017), yet the neural basis of their interaction remains unclear. Here, focusing on near– far disparity decoding, we tested how interocular suppression shapes disparity representations when neither stereopsis nor rivalry is task-relevant. Interocular suppression reduced disparity-related representations and weakened disparity-network coordination, with the strongest effects in higher-level dorsal regions. These findings provide direct neural evidence for an intrinsic coupling between binocular integration and interocular suppression, and highlight the visual dorsal stream as a key substrate for this interaction.

This dorsal bias is consistent with previous work implicating IPS in the integration of spatially distributed visual information. Studies using TMS, for example, have shown that disrupting IPS selectively alters global percept formation and illusory binding while sparing more local feature processing, consistent with a role in higher-order spatial organization (Esterman et al., 2007; Grassi et al., 2016; Zaretskaya et al., 2013). Recent 7T fMRI work further suggests a more specific role for IPS in binocular rivalry, showing enhanced eye-specific modulation in IPS together with stronger feedback from IPS to earlier visual cortex, and proposing that IPS helps bias and synchronize local eye-based competition into coherent perceptual states (Qian et al., 2026). Our findings extend this view by suggesting that IPS-related coordination is not limited to the suppressive process itself, but also shapes binocular integration across space: although rivalry was induced only in the central field, its strongest disruptive effects on disparity-related representations emerged in higher-level dorsal regions. In this framework, IPS may be a key node through which interocular suppression and binocular integration are coordinated across space.

### Anticorrelated RDS: a structured inverted code in V3A disrupted by interocular suppression

Anticorrelated random-dot stereograms (aRDS) have long provided a useful contrast for studying binocular disparity integration, because they contain disparity signals while usually failing to generate stable stereoscopic depth (Cumming & Parker, 1997). Yet under specific stimulus conditions, they can still produce a weak perceptual bias opposite to that elicited by correlated stereograms (Aoki et al., 2017; Zhaoping & Ackermann, 2018). Existing electrophysiological and neuroimaging evidence indicates that aRDS-related disparity signals can be detected across multiple stages of the visual system, although their sensitivity is transformed and often reduced relative to canonical correlated disparity representations (Alvarez et al., 2025; Cumming & Parker, 1997; Ip et al., 2014; Tanabe et al., 2004). What has remained unresolved is the representational basis of this atypical perceptual tendency. Here, a separate psychophysical test confirmed a reversed stereoscopic bias for aRDS, and cross-condition decoding combined with discriminant-axis geometry showed that in V3A this signal was expressed as a systematic inversion of the canonical correlated-disparity code. Thus, anticorrelated disparity is not merely weakened or discarded, but can be preserved in a structured, non-canonical representational format.

This inverted signature was abolished under rivalry. The significance of this result is that interocular suppression disrupted not only the canonical coding of correlated disparity, but also the structured, non-canonical representation evoked by anticorrelated stereograms. Suppression therefore appears to act at the level of binocular representational organization, rather than merely weakening the eventual depth percept. Taken together, the effects observed for both cRDS and aRDS suggest that the interaction between disparity integration and interocular suppression reflects a general organizing principle of binocular processing.

### V3A and the transformation of binocular signals into spatial representations

Prior work has established V3A as a key cortical locus for binocular disparity processing in support of stereoscopic perception (Backus et al., 2001; Ng et al., 2021; Preston et al., 2008; Tsao et al., 2003). Early fMRI work showed that stereoscopic stimulation evokes especially strong responses in V3A relative to many other retinotopic visual areas, positioning it as a major dorsal-stage site for disparity processing (Tsao et al., 2003). Subsequent studies strengthened this interpretation by showing that activity in V3A covaries closely with psychophysical limits of stereoscopic vision, including both fine stereoacuity and the upper disparity range over which depth can still be extracted (Backus et al., 2001; Chen et al., 2020). Multivoxel and ultra-high-field imaging further indicated that V3A carries perceptually relevant disparity information, contains systematic disparity-selective organization, and participates in computations that support 3D surface orientation and depth readout (Goncalves et al., 2015; Ng et al., 2021; Preston et al., 2008). Causal evidence has also converged on this view: TMS over V3A impairs fine stereoscopic judgments, indicating that V3A is not merely disparity-sensitive, but contributes directly to behaviorally relevant depth processing (Chen et al., 2020). Together, these findings have established V3A as more than an intermediate disparity-responsive area; rather, it is widely viewed as a dorsal-stage locus at which binocular disparity signals are transformed into perceptually meaningful spatial representations.

Our findings extend this account by showing that the importance of V3A is not limited to stereoscopic integration under favorable binocular conditions, but also includes its susceptibility to interocular suppression. In our data, V3A not only carried the strongest near–far disparity representation under correlated fusion, but also showed a pronounced rivalry cost under correlated rivalry, indicating that a region central to disparity-based stereopsis is also strongly affected when suppressive context is introduced. This effect was not confined to canonical disparity processing: under anticorrelated fusion, V3A expressed a structured inverted code relative to the canonical correlated-disparity format, whereas rivalry abolished this non-canonical mapping. Moreover, rivalry selectively weakened V3A-centered informational coordination across the dorsal network. Taken together, these results move the functional significance of V3A beyond its established role in stereopsis, suggesting that V3A is a key locus where binocular integration and interocular suppression converge, and where suppressive state can directly reshape both the local representational structure and the broader network organization of disparity signals.

### Clinical implications

Clinical interventions for amblyopia and strabismus often emphasize ocular endpoints, yet improvements in acuity or alignment do not necessarily translate into recovery of binocular function (Huang et al., 2007; Yu et al., 2024; Zhao et al., 2017; Zhou et al., 2017), and stereopsis often remains impaired (Feng et al., 2015; Jia et al., 2022; Xi et al., 2023). In amblyopia, stereo deficits can persist even when the amblyopic eye’s acuity returns to the normal range (Feng et al., 2015; Jia et al., 2022). In strabismus, including intermittent exotropia, good postoperative alignment likewise does not guarantee meaningful gains in stereopsis (Feng et al., 2015). These observations support the notion that rebalancing the eyes may not restore the cortical computations that sustain stereopsis (Zhou et al., 2017).

Here we identify a candidate neural bottleneck by isolating task-irrelevant interocular suppression in normal observers: suppression attenuated disparity representations most reliably in V3A and IPS and reduced coordination within a V3A-centered disparity network. These findings motivate a concrete prediction for patient studies: if suppression persists, late-dorsal disparity processing and V3A-centered network coordination may remain constrained even after ocular endpoints improve, thereby limiting stereoscopic recovery. Accordingly, V3A/IPS disparity-information readouts and V3A-centered network measures may serve as candidate neural markers for tracking intervention efficacy beyond acuity or alignment.

### Limitations and future directions

A key limitation is that our design was optimized to establish reliable effects within individuals through dense repeated measurements, rather than to characterize how these effects are distributed across the population (Schwarzkopf & Huang, 2024). Each participant completed two MRI sessions totaling approximately 5 hours of scanning, yielding high-precision within-subject estimates that enabled robust analyses of decoding performance, representational geometry, and informational connectivity. This approach is well suited for testing whether task-irrelevant interocular suppression can modulate disparity representations and their coordination in typical binocular vision. However, the small sample size limits what we can infer about individual differences in suppression-linked neural costs for disparity integration. Characterizing such variability is important because stereopsis and binocular rivalry characteristics differ markedly across individuals (Baldwin et al., 2024; Blake, 2022; Brascamp et al., 2018; Brascamp et al., 2019; Chopin et al., 2019). An important next step is therefore to broaden sampling across a wider range of stereo performance and suppression strength, and to quantify individual-difference structure in suppression effects on key neural readouts, including V3A/IPS disparity representations, V3A’s inverted aRDS mapping, and V3A-centered coupling. This can be achieved by relating these neural measures to independent behavioral and ocular metrics such as stereoacuity, ocular balance, and suppression strength.

## Materials and methods

### Participants

Eight adults with normal vision participated in the study (six females; age 22–27 years, mean ± SD = 24.1 ± 1.6 years). The sample size (N = 8) was selected to match prior fMRI decoding studies of binocular disparity (Ng et al., 2021). All participants had normal or corrected-to-normal visual acuity (logMAR ≤ 0.00), no significant anisometropia (≤ 1.00 D), astigmatism ≤ 1.50 D, and no history of ocular disease or ocular surgery. Stereovision was confirmed with a computerized random-dot stereogram test (stereo thresholds ≤ 100 arcsec). The study was approved by the Human Research Ethics Board of South China Normal University and complied with the Declaration of Helsinki. All participants provided written informed consent and received monetary compensation. Each participant completed approximately 1 hour of psychophysics testing and approximately 5 hours of MRI scanning across two sessions.

### Visual stimuli

Visual stimuli were generated in MATLAB R2022b using Psychtoolbox-3 (Brainard, 1997) and displayed on an LCD monitor (1920 × 1080 pixels, 60 Hz; 60 cm wide) viewed via a mirror at 130 cm. Dichoptic presentation was achieved using prism-based 3D glasses. The left- and right-eye images were drawn in two horizontally separated apertures on the screen, and the prisms shifted them so that they overlapped perceptually and formed a single cyclopean display. At the beginning of each session, participants used button presses to fine-tune the horizontal separation of the left- and right-eye images to achieve stable fusion. All stimuli were presented on a uniform mid-gray background.

The cyclopean display comprised three concentric regions centered on fixation (Fig. 1A). In the central fixation region, a yellow fixation dot (10-pixel diameter) was surrounded by a thin black square frame (0.5° × 0.5°) and four short white line segments aligned with the cardinal directions.

Surrounding fixation, a ring spanning 0.4°–2.0° eccentricity induced either binocular fusion or rivalry. In fusion runs, the ring contained a binocularly matched plaid generated by averaging two orthogonal sinusoidal gratings. In rivalry runs, orthogonal gratings were presented dichoptically to induce interocular suppression. Gratings had a spatial frequency of 3.75 cycles/degree; phase and orientation were randomized across blocks.

Outside 2.0° eccentricity, an 8° × 8° random-dot stereogram (RDS) aperture was filled with black-and-white dots (3-pixel diameter; density 62.5 dots/deg²). A disparity-defined annulus (2.5°–3.5° eccentricity) carried near (crossed) or far (uncrossed) horizontal disparities with three magnitudes per sign (3, 9, and 15 arcmin). Dots outside the annulus had zero disparity and were always binocularly matched. Disparity correlation was manipulated within the annulus: corresponding dots had the same contrast polarity in correlated conditions (black–black or white–white) but opposite polarity in anticorrelated conditions (black–white). The RDS aperture was embedded within a 9° × 9° high-contrast checkerboard surround that served as a fusion frame to facilitate stable binocular alignment.

### Experimental design and tasks

**Pre-scan stereo discrimination task.** Before scanning, participants completed a stereo discrimination task to assess near–far judgments from the disparity-defined annulus. The stimulus configuration matched that used in the fMRI experiment, crossing disparity correlation (correlated, anticorrelated) with interocular state (fusion, rivalry). On each trial, the annulus carried one of 12 horizontal disparities (±1.5, ±2.25, ±3, ±4.5, ±9, ±15 arcmin). After 20 practice trials with trial-wise feedback, participants completed 960 trials (12 disparities × 4 conditions × 20 repetitions), divided into eight blocks of 120 trials with self-paced breaks. Each trial consisted of a 0.5-s fixation-only interval, followed by a 1.0-s stimulus interval and a 1.0-s response interval. Participants reported whether the annulus appeared nearer or farther than fixation using the up/down arrow keys while maintaining central fixation.

**fMRI main experiment.** The fMRI experiment was conducted across two sessions on separate days. Each session included 16 runs of the main binocular disparity experiment, one annulus localizer run, and a high-resolution T1-weighted anatomical scan. In one session, participants additionally completed six 5-min runs of a standard population receptive field (pRF) mapping protocol (Benson & Winawer, 2018).

In the main experiment, each run contained a single condition defined by crossing disparity correlation (correlated vs anticorrelated) and interocular state (fusion vs rivalry): correlated fusion (CF), correlated rivalry (CR), anticorrelated fusion (AF), and anticorrelated rivalry (AR). Across 32 runs, conditions were balanced across sessions and participants (8 runs per condition), with run order counterbalanced across sessions.

Each main run comprised 151 volumes (TR = 2 s; total duration 302 s) and consisted of 18 blocks. Within each block, the disparity-defined annulus was presented in 10 stimulus cycles (0.5 s blank followed by 1.0 s stimulus), yielding a 15-s block duration. Near and far blocks were balanced within each run (9 near, 9 far). Disparity magnitude (3, 9, 15 arcmin) was randomized across blocks within each disparity sign. Across the full dataset, this design yielded 144 blocks per condition (72 near, 72 far), which served as inputs to the decoding analyses.

**Fixation-change detection task.** To minimize top–down influences, participants performed a fixation-change detection task throughout scanning. Fixation-change events occurred 60 times per run and were pseudo-randomly distributed. On each event, the fixation dot transiently changed into a horizontally elongated ellipse (16 × 10 pixels) or a vertically elongated ellipse (10 × 16 pixels) for 300, 400, or 500 ms, while the surrounding frame and line segments remained unchanged. Participants indicated the direction of elongation using two buttons on an MR-compatible response box and were instructed to ignore perceived depth and any dominance changes in the surrounding stimuli.

### MRI data acquisition

MRI data were acquired on a 3T Siemens MAGNETOM Prisma scanner with a 32-channel head coil. Functional images were collected using a T2*-weighted gradient-echo EPI sequence (TR = 2000 ms, TE = 29 ms, flip angle = 90°, voxel size = 1.5 × 1.5 × 2.0 mm³, field of view = 180 mm, matrix = 120 × 120, 30 slices, slice thickness = 2.0 mm, no slice gap). Slices were acquired in interleaved order with a near-coronal prescription tilted anteriorly to optimize coverage of occipital and dorsal visual areas. Phase encoding was along the foot–head direction. Parallel imaging was applied (GRAPPA acceleration factor = 2) to shorten EPI readout and reduce susceptibility-related distortions.

In each session, an annulus localizer run was acquired with the same EPI parameters (196 volumes). In one session, participants additionally completed six pRF mapping runs acquired with the same EPI parameters (150 volumes per run).

High-resolution anatomical images were acquired using a T1-weighted 3D MPRAGE sequence (TR = 2530 ms, TE = 2.27 ms, TI = 1100 ms, flip angle = 7°, voxel size = 1.0 × 1.0 × 1.0 mm³, field of view = 256 × 256 mm², matrix = 256 × 256, 208 sagittal slices, no slice gap) with in-plane GRAPPA acceleration (factor = 2).

### MRI preprocessing

T1-weighted anatomical images from the two sessions were processed with FreeSurfer (Fischl, 2012) to reconstruct individual cortical surfaces. Functional data were preprocessed in AFNI (Cox, 1996) using a consistent pipeline across the main experiment, the annulus localizer, and the pRF runs. Preprocessing included slice-timing correction, rigid-body motion correction, and EPI-to-T1 registration; alignment quality was verified by visual inspection. No spatial smoothing was applied to preserve fine-grained multivariate patterns.

### Block-wise response estimation for the main experiment (LSS)

Block-wise responses in the main experiment were estimated using a least-squares–separate (LSS) GLM (Mumford et al., 2012) implemented in AFNI. For each block, we fit a model with a regressor for the target block and separate regressors for all remaining blocks, with six motion parameters and third-order polynomial trends included as nuisance regressors. This procedure yielded one β map per block. Across 32 runs (18 blocks/run), we obtained 576 β maps (144 per condition; balanced near/far) for subsequent decoding analyses.

### ROI definition

Subject-specific retinotopic ROIs (V1, V2, V3, V3A, hV4, and VO) were defined from pRF mapping. For the pRF runs, preprocessed time series were resampled to TR = 1 s, mean-centered, and detrended with a third-order polynomial. Population receptive-field parameters were estimated using the qPRF toolbox (Waz et al., 2025), and neuropythy (Benson & Winawer, 2018) was used for Bayesian retinotopic mapping to derive visual area labels on the cortical surface. These labels were then converted to native-volume ROI masks for MVPA. The VO ROI was defined by combining VO1 and VO2 into a single ventral occipital ROI.

To define IPS, we generated an atlas-based parcellation using the Wang et al. (2015) atlas and combined IPS0–IPS5 into a single IPS ROI.

Because decoding targeted the disparity-defined annulus (2.5°–3.5° eccentricity), voxels in V1– V3A were further restricted using an independent annulus localizer (annulus > fixation, p < 0.05, uncorrected). IPS was not constrained by the localizer, as its responses are less strictly retinotopic.

### Multivariate pattern analysis

Multivariate analyses were implemented in Python using scikit-learn. Voxel-wise β values were z-scored within each run before all multivariate analyses.

**Within-condition decoding.** Within each condition (CF, CR, AF, AR) and ROI, we trained a linear support vector machine (SVM; scikit-learn LinearSVC, C = 1.0) to discriminate near versus far disparity from voxel-wise β patterns. Decoding performance was evaluated using leave-one-run-out cross-validation within each condition. Each condition comprised 8 runs (18 blocks/run; 144 blocks total). In each fold, one run was held out for testing and the classifier was trained on the remaining seven runs. Accuracy was computed on the held-out run and averaged across folds, yielding one decoding accuracy per participant × condition × ROI (chance level = 0.5).

**Cross-condition decoding.** To assess generalization of disparity representations across conditions, we performed cross-condition decoding for selected condition pairs. For each ROI, an SVM was trained on all runs from condition A and tested on all runs from condition B. The analysis was repeated in the reverse direction (train on B, test on A), and the two accuracies were averaged to obtain a symmetric cross-decoding metric:

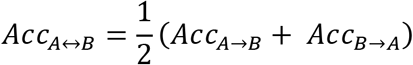

As a within-condition reference, we computed split-half decoding within correlated fusion (CF) by treating odd and even CF runs as two independent partitions (A and B) and applying the same bidirectional train–test procedure as in cross-condition decoding to yield a symmetric split-half CF metric.

**Decision-axis angle analysis.** To quantify the similarity of near–far coding across conditions within each ROI, we compared classifier decision axes between conditions. Using the same linear SVM settings as in the decoding analyses, we fit a classifier for each participant, ROI, and condition using all blocks from that condition and extracted the SVM weight vector *w* as the decision axis. The angle between two axes was computed as:

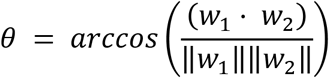

yielding *θ* ∈ [0°, 180°]. We summarized deviation from orthogonality as:

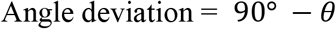

where positive values indicate more similar axes and negative values indicate inverted axes.

Axis reliability in correlated fusion (CF) was assessed with a split-half procedure across runs: the classifier axis was estimated separately using odd-numbered CF runs and even-numbered CF runs.

**Informational connectivity.** We quantified cross-ROI coupling of disparity information using informational connectivity. For each ROI and condition, we obtained a cross-validated decision-value series using leave-one-run-out decoding: in each fold, a classifier trained on seven runs was applied to the held-out run to yield signed decision values (distance to the hyperplane) for each block. Decision values were concatenated across folds to form a cross-validated series (144 blocks) per ROI and condition for each participant. Informational connectivity between a pair of ROIs was defined as the Pearson correlation between their decision-value series within the same condition, with correlation coefficients Fisher z-transformed before group-level averaging and statistical testing. In each participant and condition, we focused on two network summaries: mean connectivity within early visual cortex (V1–V3) and mean connectivity within a V3A-centered network (all pairs involving V3A).

### Statistical analysis

To test whether the rivalry cost for correlated disparity decoding increased along the dorsal visual hierarchy, we fitted a linear mixed-effects model to the CF–CR decoding difference across V1, V2, V3, V3A, and IPS, with ROI level as an ordered fixed effect and subject as a random intercept. Normality of participant-level estimates was assessed with Shapiro–Wilk tests. One-sided tests were used for within-condition decoding above chance, informational connectivity above zero, and rivalry cost in the correlated condition (CF − CR); all other tests were two-sided. Multiple comparisons were controlled using the Holm–Bonferroni method within each analysis family, with the corresponding correction schemes detailed in the table notes. For clarity, we report only key statistics in the main text; complete results are provided in the Supplementary materials (Tables S1–S7).

## Data and code availability

Behavioral data, MVPA-derived files (ROI masks/labels and block-wise beta estimates), and all analysis and figure-generation scripts are available on Zenodo (DOI: 10.5281/zenodo.17958394).

## Acknowledgments

Supported by grants from the STI2030-Major Projects (2021ZD0204200), the National Natural Science Foundation of China (32371100), and the Research Center for Brain Cognition and Human Development, Guangdong, China (2024B0303390003).

## Supplementary materials

**Table S1.** Psychophysical depth-discrimination performance.

| Comparison | Mean ± SD | *t* | df | *p*_Holm | Cohen's *d* |
| --- | --- | --- | --- | --- | --- |
| CF vs chance | 0.958 ± 0.066 | 19.67 | 7 | <0.001 | 6.95 |
| CR vs chance | 0.970 ± 0.038 | 35.42 | 7 | <0.001 | 12.52 |
| AF vs chance | 0.410 ± 0.061 | -4.17 | 7 | 0.004 | -1.48 |
| AR vs chance | 0.399 ± 0.042 | -6.82 | 7 | <0.001 | -2.41 |
| CF vs CR | -0.011 ± 0.038 | -0.86 | 7 | 0.839 | -0.30 |
| CF vs CR | 0.011 ± 0.063 | 0.51 | 7 | 0.839 | 0.18 |

**Note.** All tests were two-sided *t*-tests. CF = correlated fusion; CR = correlated rivalry; AF = anticorrelated fusion; AR = anticorrelated rivalry.

**Table S2.** Within-condition decoding accuracy above chance.

| Category | Condition | ROI | Mean ± SD | *t* | df | *p*_Holm | Cohen's *d* |
| --- | --- | --- | --- | --- | --- | --- | --- |
| Early | CF | V1 | 0.558 ± 0.041 | 3.97 | 7 | 0.008 | 1.40 |
| Early | CF | V2 | 0.583 ± 0.064 | 3.67 | 7 | 0.008 | 1.30 |
| Early | CF | V3 | 0.606 ± 0.086 | 3.46 | 7 | 0.008 | 1.22 |
| Early | CR | V1 | 0.544 ± 0.061 | 2.05 | 7 | 0.062 | 0.73 |
| Early | CR | V2 | 0.549 ± 0.062 | 2.21 | 7 | 0.062 | 0.78 |
| Early | CR | V3 | 0.555 ± 0.056 | 2.75 | 7 | 0.043 | 0.97 |
| Early | AF | V1 | 0.498 ± 0.045 | -0.11 | 7 | 0.542 | -0.04 |
| Early | AF | V2 | 0.517 ± 0.050 | 0.98 | 7 | 0.361 | 0.35 |
| Early | AF | V3 | 0.524 ± 0.051 | 1.36 | 7 | 0.325 | 0.48 |
| Early | AR | V1 | 0.492 ± 0.044 | -0.51 | 7 | 1.000 | -0.18 |
| Early | AR | V2 | 0.511 ± 0.076 | 0.42 | 7 | 1.000 | 0.15 |
| Early | AR | V3 | 0.482 ± 0.079 | -0.65 | 7 | 1.000 | -0.23 |
| Dorsal | CF | V3A | 0.673 ± 0.071 | 6.86 | 7 | <0.001 | 2.42 |
| Dorsal | CF | IPS | 0.605 ± 0.068 | 4.36 | 7 | 0.002 | 1.54 |
| Dorsal | CR | V3A | 0.597 ± 0.058 | 4.70 | 7 | 0.002 | 1.66 |
| Dorsal | CR | IPS | 0.509 ± 0.045 | 0.54 | 7 | 0.302 | 0.19 |
| Dorsal | AF | V3A | 0.517 ± 0.063 | 0.78 | 7 | 0.463 | 0.27 |
| Dorsal | AF | IPS | 0.489 ± 0.062 | -0.52 | 7 | 0.689 | -0.18 |
| Dorsal | AR | V3A | 0.515 ± 0.063 | 0.67 | 7 | 0.481 | 0.24 |
| Dorsal | AR | IPS | 0.516 ± 0.059 | 0.74 | 7 | 0.481 | 0.26 |
| Ventral | CF | hV4 | 0.549 ± 0.069 | 2.03 | 7 | 0.041 | 0.72 |
| Ventral | CF | VO | 0.571 ± 0.065 | 3.08 | 7 | 0.018 | 1.09 |
| Ventral | CR | hV4 | 0.538 ± 0.058 | 1.86 | 7 | 0.064 | 0.66 |
| Ventral | CR | VO | 0.537 ± 0.048 | 2.20 | 7 | 0.064 | 0.78 |
| Ventral | AF | hV4 | 0.505 ± 0.055 | 0.27 | 7 | 0.399 | 0.09 |
| Ventral | AF | VO | 0.530 ± 0.044 | 1.90 | 7 | 0.100 | 0.67 |
| Ventral | AR | hV4 | 0.497 ± 0.055 | -0.13 | 7 | 0.650 | -0.05 |
| Ventral | AR | VO | 0.507 ± 0.042 | 0.47 | 7 | 0.650 | 0.17 |

**Note.** All tests were one-sided one-sample *t*-tests against chance, and *p* values were corrected using the Holm–Bonferroni method within each ROI category. CF = correlated fusion; CR = correlated rivalry; AF = anticorrelated fusion; AR = anticorrelated rivalry; ROI = region of interest.

**Table S3.** Rivalry cost in within-condition decoding.

| Category | Condition | ROI | Mean_diff ± SD | *t* | df | *p*_Holm | Cohen's *d* |
| --- | --- | --- | --- | --- | --- | --- | --- |
| Early | Correlated | V1 | 0.014 ± 0.094 | 0.42 | 7 | 0.344 | 0.15 |
| Early | Correlated | V2 | 0.035 ± 0.042 | 2.32 | 7 | 0.053 | 0.82 |
| Early | Correlated | V3 | 0.051 ± 0.047 | 3.09 | 7 | 0.027 | 1.09 |
| Early | Anticorrelated | V1 | 0.006 ± 0.045 | 0.39 | 7 | 1.000 | 0.14 |
| Early | Anticorrelated | V2 | 0.006 ± 0.106 | 0.16 | 7 | 1.000 | 0.06 |
| Early | Anticorrelated | V3 | 0.043 ± 0.103 | 1.17 | 7 | 0.842 | 0.41 |
| Dorsal | Correlated | V3A | 0.076 ± 0.077 | 2.76 | 7 | 0.014 | 0.98 |
| Dorsal | Correlated | IPS | 0.096 ± 0.066 | 4.15 | 7 | 0.004 | 1.47 |
| Dorsal | Anticorrelated | V3A | 0.003 ± 0.096 | 0.08 | 7 | 0.941 | 0.03 |
| Dorsal | Anticorrelated | IPS | -0.027 ± 0.086 | -0.88 | 7 | 0.812 | -0.31 |
| Ventral | Correlated | hV4 | 0.011 ± 0.067 | 0.48 | 7 | 0.324 | 0.17 |
| Ventral | Correlated | VO | 0.034 ± 0.083 | 1.15 | 7 | 0.286 | 0.41 |
| Ventral | Anticorrelated | hV4 | 0.008 ± 0.088 | 0.25 | 7 | 0.809 | 0.09 |
| Ventral | Anticorrelated | VO | 0.023 ± 0.045 | 1.42 | 7 | 0.400 | 0.50 |

**Note.** Mean\_diff refers to fusion minus rivalry within each condition. Correlated effects (CF – CR) were tested one-sided, whereas anticorrelated effects (AF – AR) were tested two-sided. *p* values were corrected using the Holm–Bonferroni method within each ROI category. CF = correlated fusion; CR = correlated rivalry; AF = anticorrelated fusion; AR = anticorrelated rivalry; ROI = region of interest.

**Table S4.** Cross-condition decoding accuracy relative to chance level.

| Category | Comparison | ROI | Mean ± SD | *t* | df | *p*<sub>Holm</sub> | Cohen's *d* |
| --- | --- | --- | --- | --- | --- | --- | --- |
| Early | CF split-half | V1 | 0.583 ± 0.055 | 4.32 | 7 | 0.003 | 1.53 |
| Early | CF split-half | V2 | 0.619 ± 0.038 | 8.87 | 7 | <0.001 | 3.13 |
| Early | CF split-half | V3 | 0.618 ± 0.029 | 11.33 | 7 | <0.001 | 4.01 |
| Early | CF ↔ CR | V1 | 0.536 ± 0.034 | 3.05 | 7 | 0.037 | 1.08 |
| Early | CF ↔ CR | V2 | 0.574 ± 0.042 | 4.95 | 7 | 0.005 | 1.75 |
| Early | CF ↔ CR | V3 | 0.566 ± 0.062 | 3.02 | 7 | 0.037 | 1.07 |
| Early | CF ↔ AF | V1 | 0.498 ± 0.025 | -0.24 | 7 | 1.000 | -0.09 |
| Early | CF ↔ AF | V2 | 0.497 ± 0.042 | -0.17 | 7 | 1.000 | -0.06 |
| Early | CF ↔ AF | V3 | 0.482 ± 0.047 | -1.08 | 7 | 0.949 | -0.38 |
| Early | CF ↔ AR | V1 | 0.486 ± 0.022 | -1.84 | 7 | 0.324 | -0.65 |
| Early | CF ↔ AR | V2 | 0.487 ± 0.030 | -1.26 | 7 | 0.495 | -0.45 |
| Early | CF ↔ AR | V3 | 0.492 ± 0.020 | -1.10 | 7 | 0.495 | -0.39 |
| Dorsal | CF split-half | V3A | 0.651 ± 0.081 | 5.24 | 7 | 0.001 | 1.85 |
| Dorsal | CF split-half | IPS | 0.621 ± 0.040 | 8.43 | 7 | <0.001 | 2.98 |
| Dorsal | CF ↔ CR | V3A | 0.608 ± 0.064 | 4.75 | 7 | 0.004 | 1.68 |
| Dorsal | CF ↔ CR | IPS | 0.583 ± 0.062 | 3.83 | 7 | 0.006 | 1.35 |
| Dorsal | CF ↔ AF | V3A | 0.461 ± 0.037 | -3.01 | 7 | 0.039 | -1.06 |
| Dorsal | CF ↔ AF | IPS | 0.484 ± 0.045 | -1.04 | 7 | 0.334 | -0.37 |
| Dorsal | CF ↔ AR | V3A | 0.495 ± 0.069 | -0.21 | 7 | 1.000 | -0.08 |
| Dorsal | CF ↔ AR | IPS | 0.501 ± 0.033 | 0.07 | 7 | 1.000 | 0.03 |

**Note.** All tests were two-sided one-sample *t*-tests against chance, and *p* values were corrected using the Holm–Bonferroni method within each ROI category separately for each comparison. CF = correlated fusion; CR = correlated rivalry; AF = anticorrelated fusion; AR = anticorrelated rivalry; ROI = region of interest.

**Table S5.** Cross-condition axis-angle deviation from orthogonality.

| Comparison | ROI | Mean ± SD | *t* | df | *p*_Holm | Cohen's *d* |
| --- | --- | --- | --- | --- | --- | --- |
| CF split-half | V3A | 5.78 ± 3.76 | 4.34 | 7 | 0.004 | 1.54 |
| CF split-half | IPS | 2.49 ± 1.47 | 4.80 | 7 | 0.004 | 1.70 |
| CF vs CR | V3A | 5.90 ± 2.88 | 5.79 | 7 | 0.001 | 2.05 |
| CF vs CR | IPS | 3.14 ± 2.70 | 3.29 | 7 | 0.013 | 1.16 |
| CF vs AF | V3A | -2.21 ± 1.96 | -3.19 | 7 | 0.031 | -1.13 |
| CF vs AF | IPS | -0.73 ± 2.34 | -0.89 | 7 | 0.405 | -0.31 |
| CF vs AR | V3A | -0.46 ± 4.33 | -0.30 | 7 | 0.775 | -0.11 |
| CF vs AR | IPS | 0.35 ± 0.99 | 0.99 | 7 | 0.715 | 0.35 |

**Note.** Axis-angle deviation values are reported as deviations from 90° (i.e., orthogonality), such that positive values indicate angles smaller than 90° and negative values indicate angles larger than 90°. All tests were two-sided one-sample *t*-tests against zero. For this dorsal-stream table, *p* values were corrected using the Holm–Bonferroni method across ROIs within each comparison. CF = correlated fusion; CR = correlated rivalry; AF = anticorrelated fusion; AR = anticorrelated rivalry; ROI = region of interest.

**Table S6.** Informational connectivity within each condition (Fisher z)

| Condition | Category | ROI pairs | Mean ± SD | *t* | df | *p*_Holm | Cohen's *d* |
| --- | --- | --- | --- | --- | --- | --- | --- |
| CF | Early | V1-V2 | 0.187 ± 0.104 | 5.08 | 7 | 0.002 | 1.80 |
| CF | Early | V2-V3 | 0.229 ± 0.160 | 4.04 | 7 | 0.005 | 1.43 |
| CF | Early | V1-V3 | 0.166 ± 0.138 | 3.40 | 7 | 0.006 | 1.20 |
| CF | V3A-centered | V3A-V1 | 0.180 ± 0.134 | 3.80 | 7 | 0.005 | 1.34 |
| CF | V3A-centered | V3A-V2 | 0.215 ± 0.150 | 4.05 | 7 | 0.005 | 1.43 |
| CF | V3A-centered | V3A-V3 | 0.317 ± 0.189 | 4.74 | 7 | 0.003 | 1.68 |
| CF | V3A-centered | V3A-IPS | 0.374 ± 0.179 | 5.89 | 7 | 0.001 | 2.08 |
| CR | Early | V1-V2 | 0.165 ± 0.086 | 5.40 | 7 | 0.001 | 1.91 |
| CR | Early | V2-V3 | 0.235 ± 0.161 | 4.13 | 7 | 0.002 | 1.46 |
| CR | Early | V1-V3 | 0.137 ± 0.060 | 6.47 | 7 | 0.001 | 2.29 |
| CR | V3A-centered | V3A-V1 | 0.048 ± 0.066 | 2.05 | 7 | 0.040 | 0.72 |
| CR | V3A-centered | V3A-V2 | 0.104 ± 0.078 | 3.74 | 7 | 0.007 | 1.32 |
| CR | V3A-centered | V3A-V3 | 0.185 ± 0.129 | 4.08 | 7 | 0.007 | 1.44 |
| CR | V3A-centered | V3A-IPS | 0.271 ± 0.096 | 8.00 | 7 | <0.001 | 2.83 |
| AF | Early | V1-V2 | 0.095 ± 0.212 | 1.27 | 7 | 0.245 | 0.45 |
| AF | Early | V2-V3 | 0.216 ± 0.122 | 5.02 | 7 | 0.002 | 1.77 |
| AF | Early | V1-V3 | 0.023 ± 0.116 | 0.56 | 7 | 0.297 | 0.20 |
| AF | V3A-centered | V3A-V1 | 0.071 ± 0.065 | 3.12 | 7 | 0.033 | 1.11 |
| AF | V3A-centered | V3A-V2 | 0.040 ± 0.102 | 1.12 | 7 | 0.150 | 0.40 |
| AF | V3A-centered | V3A-V3 | 0.062 ± 0.060 | 2.92 | 7 | 0.034 | 1.03 |
| AF | V3A-centered | V3A-IPS | 0.100 ± 0.135 | 2.10 | 7 | 0.074 | 0.74 |
| AR | Early | V1-V2 | 0.113 ± 0.170 | 1.88 | 7 | 0.051 | 0.67 |
| AR | Early | V2-V3 | 0.094 ± 0.096 | 2.79 | 7 | 0.040 | 0.99 |
| AR | Early | V1-V3 | 0.112 ± 0.125 | 2.54 | 7 | 0.040 | 0.90 |
| AR | V3A-centered | V3A-V1 | 0.016 ± 0.072 | 0.65 | 7 | 0.270 | 0.23 |
| AR | V3A-centered | V3A-V2 | 0.082 ± 0.118 | 1.96 | 7 | 0.091 | 0.69 |
| AR | V3A-centered | V3A-V3 | 0.108 ± 0.058 | 5.22 | 7 | 0.002 | 1.85 |
| AR | V3A-centered | V3A-IPS | 0.178 ± 0.095 | 5.30 | 7 | 0.002 | 1.87 |

**Note.** All tests were one-sided one-sample *t*-tests against zero, and *p* values were corrected using the Holm–Bonferroni method within each ROI category separately for each condition. Early ROI pairs included V1-V2, V2-V3, and V1-V3; V3A-centered ROI pairs included V3A-V1, V3A-V2,
V3A-V3, and V3A-IPS. CF = correlated fusion; CR = correlated rivalry; AF = anticorrelated fusion; AR = anticorrelated rivalry; ROI = region of interest.

**Table S7.** Informational connectivity contrasts (Fisher z) across conditions.

| Condition | ROI pair | Mean ± SD | *t* | df | *p*<sub>Holm</sub> | Cohen's *d* |
| --- | --- | --- | --- | --- | --- | --- |
| Correlated | V1-V2 | 0.022 ± 0.111 | 0.57 | 7 | 1.000 | 0.20 |
| Correlated | V2-V3 | -0.007 ± 0.267 | -0.07 | 7 | 1.000 | -0.03 |
| Correlated | V1-V3 | 0.028 ± 0.142 | 0.57 | 7 | 1.000 | 0.20 |
| Correlated | Early mean | 0.015 ± 0.142 | 0.29 | 7 | 1.000 | 0.10 |
| Correlated | V3A-V1 | 0.133 ± 0.128 | 2.93 | 7 | 0.044 | 1.03 |
| Correlated | V3A-V2 | 0.111 ± 0.166 | 1.89 | 7 | 0.100 | 0.67 |
| Correlated | V3A-V3 | 0.132 ± 0.136 | 2.75 | 7 | 0.044 | 0.97 |
| Correlated | V3A-IPS | 0.103 ± 0.205 | 1.43 | 7 | 0.100 | 0.50 |
| Correlated | V3A-centered mean | 0.120 ± 0.096 | 3.52 | 7 | 0.024 | 1.24 |
| Anticorrelated | V1-V2 | -0.018 ± 0.157 | -0.33 | 7 | 1.000 | -0.12 |
| Anticorrelated | V2-V3 | 0.122 ± 0.134 | 2.57 | 7 | 0.148 | 0.91 |
| Anticorrelated | V1-V3 | -0.089 ± 0.143 | -1.76 | 7 | 0.363 | -0.62 |
| Anticorrelated | Early mean | 0.005 ± 0.119 | 0.11 | 7 | 1.000 | 0.04 |
| Anticorrelated | V3A-V1 | 0.055 ± 0.095 | 1.64 | 7 | 0.656 | 0.58 |
| Anticorrelated | V3A-V2 | -0.042 ± 0.075 | -1.58 | 7 | 0.656 | -0.56 |
| Anticorrelated | V3A-V3 | -0.046 ± 0.078 | -1.66 | 7 | 0.656 | -0.59 |
| Anticorrelated | V3A-IPS | -0.078 ± 0.163 | -1.35 | 7 | 0.656 | -0.48 |
| Anticorrelated | V3A-centered mean | -0.028 ± 0.046 | -1.71 | 7 | 0.656 | -0.60 |

**Note.** Mean values indicate connectivity differences between conditions (CF – CR for the correlated condition and AF – AR for the anticorrelated condition). Correlated differences were assessed using one-sided paired-sample *t*-tests, whereas anticorrelated differences were assessed using two-sided paired-sample *t*-tests. *p* values were corrected using the Holm–Bonferroni method within each ROI category. Early ROI pairs included V1-V2, V2-V3, and V1-V3; V3A-centered ROI pairs included V3A-V1, V3A-V2, V3A-V3, and V3A-IPS. CF = correlated fusion; CR = correlated rivalry; AF = anticorrelated fusion; AR = anticorrelated rivalry; ROI = region of interest.

## Notes

### Competing Interest Statement

The authors have declared no competing interest.

